# Integrating zoonotic pathogen spillover risk into Life Cycle Assessment: Probabilistic method development for chicken farming

**DOI:** 10.1101/2025.10.18.683192

**Authors:** John D. Hader, Nadia Malinverno, Claudia Som

## Abstract

Animal agriculture contributes substantially to the risk of emerging zoonotic diseases, which are an environmental and human health risk not captured by current life cycle assessment (LCA) methods for animal products. Furthermore, new value chains in the bioeconomy are emerging to valorize sidestreams from the farmed animal industry, such as using feathers for protein-based materials, potentially leading to changing pathogen exposures. Methods for assessing the risk of zoonotic pathogen spillovers across the life cycle of animal agriculture are thus critically needed. We present a conceptual framework for the life cycle risk of zoonotic pathogen spillovers in industrial-scale chicken production, based around the spillover risk at each stage of chicken production (i.e., breeding, hatching, grow-out, processing, and sidestream valorization). We develop an initial semi-quantification for the spillover risk of avian influenza at the grow-out stage in two key chicken producing countries (Brazil and the United States) utilizing event tree modelling. Our preliminary findings suggest spillover risk at the grow-out stage is higher in the US than Brazil, owing in part to a larger presence of migratory bird species (key reservoirs of avian influenza). Additional research is necessary to quantify risk distributions for other production stages and develop methods for allocating spillover risks (especially to emerging valorization of sidestreams). Incorporating the risk of zoonotic pathogen spillovers into animal product LCA – as has been done for e.g., greenhouse gas emissions and land use – would help inform policymakers and consumers regarding the impacts of dietary choices and contribute to international One Health goals.

## 1. Introduction

Global demand for animal-derived meat, eggs, and dairy products is at historical highs (Sans and Combris, 2015). Roughly 2.7E+10 chickens, 9.7E+8 pigs, and 1.6E+9 cattle are farmed globally at a given time (FAOSTAT, 2025). The number of farmed chickens, for example, is over an order of magnitude larger than the number of birds in the largest wild bird species (Bennett et al., 2018). Breeding, raising, feeding, killing, and processing this number of animals has a large impact on the environment, with animal agriculture accounting for roughly 20% of greenhouse gas (GHG) emissions globally, 40% of habitable land use, and making up the majority of antibiotic usage globally (Agudelo Higuita et al., 2023; Mulchandani et al., 2023; Van Boeckel et al., 2017). Furthermore, processing of slaughtered animals produces substantial amounts of sidestreams, such as bones, blood, and feathers (Peydayesh and Bieri, 2025), with roughly 47 Mt of animal byproducts or waste produced at the processing and manufacturing stage annually in Europe alone (Caldeira et al., 2019).

In addition to its negative environmental impacts, animal agriculture also plays a large role in emerging zoonotic diseases – i.e., diseases caused by pathogens that transmit from non-human animals to humans (Carlson et al., 2025; Hayek, 2022; Jones et al., 2013; Rohr et al., 2019; UNEP, 2016). Such emerging infectious diseases largely circulate in wildlife, and while they can pose a direct risk to humans from wildlife, they are often limited in their transmissibility from wild animals to humans. However, farmed animals can act as intermediary species between wildlife and humans for emerging diseases (Carlson et al., 2025; UNEP, 2016). Farmed animals’ potential contact with wildlife, large numbers, low genetic diversity, and often unhealthy conditions increase the possibility for infection and mutations or mixing of virus strains. This can lead to either a more efficient route of the pathogen from wild animals into humans, and/or variants of the infectious disease emerging that can more easily infect (or “spill over”) into humans (Bartlett et al., 2022; Carlson et al., 2025; Linder et al., 2023; UNEP, 2016).

A recent report on zoonotic disease risk in United States (US) animal markets highlighted that various practices in animal agriculture have differential impact on zoonotic disease risk (Linder et al., 2023). The number, type, and manner of how animals are housed, genetic diversity of animals, whether multiple species are held in close proximity to each other, types of pathogens the animals are susceptible to, the overall health of the animals, their interactions with humans during the production process, possibilities for interaction with wildlife, supply chain length/method of transport, and scale of the industry all impact the risks in the consumer markets of animals. Animal farming can also act as an amplifier of the pathogenicity of viruses, which can then “spill” back into wildlife (Carlson et al., 2025; Jori et al., 2021). Various studies have reported on the risk factors for zoonotic disease associated with farmed animals and have identified potential hot spots for zoonotic disease risk. For example, (Thanapongtharm et al., 2019) analyzed pig farming locations in Thailand, and compared these to locations and activity regions of colonies of bats (carriers of Nipah virus) to understand where the greatest likelihood of spillover from wildlife into farmed animals (and thus humans) might be. Furthermore, both industrial-scale and small-scale animal agriculture operations pose risks for zoonotic disease emergence, amplification, and spread (e.g., (Bartlett et al., 2022)).

Increasing human population and rising incomes in some regions globally are expected to increase global demand for animal products in the coming decades. By 2050, demand for meat is expected to increase 75% relative to 2005 (Alexandratos and Bruinsma, 2012). Despite these projections, the impact of animal agriculture on GHG emissions, land use, and other environmental factors (as well as the negative health effects of red and processed meat) are gaining public awareness and are increasingly being discussed in national and international documents in regards to increasing the sustainability and healthfulness of food systems (European Commission, 2023). However, zoonotic disease risks from animal agriculture are an often omitted aspect of these discussions (Verkuijl et al., 2024), and are frequently left out of quantitative assessments of animal agriculture due to lack of data (Arrigoni et al., 2023; Sinke et al., 2023). Furthermore, proposals by the Food and Agriculture Organization of the United Nations (FAO) for alleviating some of these environmental concerns (e.g., shifting from beef to more chicken production due to chicken’s lower environmental footprint; see (Poore and Nemecek, 2018)) could result in increased impacts on poorly assessed aspects like zoonotic disease risk (Verkuijl et al., 2024).

Additionally, various emerging methods within the bioeconomy have explored ways of valorizing the processing sidestreams from animal products into potentially more sustainable and value-added products, such as extracting keratin from chicken feathers for use in advanced protein-based materials; (Soon et al., 2023). With these sustainability-oriented emerging technologies, it is crucial to allocate the background sustainability impacts of the original product to these sidestreams if valorized in other methods (including any zoonotic risks). Additionally, changing methods for valorization of animal product sidestreams could also change how humans interact with these sidestreams relative to current valorization methods (e.g., landfilling or incineration; (Tesfaye et al., 2017)), and thus potentially impact zoonotic pathogen spillover risks associated with these processing sidestreams.

With this overall changing landscape of farmed animal production and demand, and potential shifts in the use of animal-derived waste streams, incorporation of zoonotic disease risks from animal agriculture into sustainability assessments would help guide decision making and provide a more complete understanding of the impacts of animal agriculture on the environment and humans. Quantifying, or at least semi-quantifying, the zoonotic pathogen spillover risks at each stage of the animal agriculture value chain, and under different production methods, would enable comparison of the zoonotic disease risks from different practices and animal products, and provide increased transparency to the overall environmental impact of different agricultural products. To the best of the authors’ knowledge, no analysis has quantitatively investigated how the activities along each step of the value chain (i.e., the life cycle) of different animal farming practices contribute to an increased risk of zoonotic disease spillover. In this study, we aim to begin filling the gap around the risk of zoonotic disease in sustainability assessments of animal agriculture (and the bioeconomy it could support) by:

- Providing an **overview of industrial chicken production**, and areas within this that are risk factors for zoonotic pathogen spillover.
- Developing a **framework for incorporating the risk of zoonotic pathogen spillover into life cycle assessment (LCA) of chicken farming**, providing initial semi-quantification of risks for one of the key stages of chicken production (grow-out).
- Outline **key knowledge and data gaps** that need to be filled to more fully capture this environmental impact of animal agricultural practices.

We limit the scope of this study to industrial chicken farming due to the rapidly growing demand for chicken meat globally (Bennett et al., 2018), the large number of farmed chickens globally compared to other animals like pigs and cattle (FAOSTAT, 2025), and their capacity for being reservoirs and transmitters of zoonotic pathogens (Carlson et al., 2025). Furthermore, we will focus our analysis on a zoonotic disease caused by a virus prioritized as being a virus with pandemic potential, specifically highly pathogenic avian influenza (HPAI) (Opata et al., 2025; Possas et al., 2025; WHO, 2024). Other pathogens associated with animal agriculture like those in feces or pathogens passed from animals to humans via insects also pose a risk to human health (Carlson et al., 2025; Linder et al., 2023; Rohr et al., 2019), but are not considered in this study. Additionally, agricultural practices other than just the farming of animals pose some risk for the emergence or spread of disease, such as land clearing for human-consumed crops (Morand and Lajaunie, 2021). However, since animal agriculture is a massive contributor to land use (for both feed crop production and grazing land; (Agudelo Higuita et al., 2023)) and deforestation (Pendrill et al., 2019), and since there are unique concerns around agricultural animals being reservoirs, amplifiers, and transmitters to humans of pandemic-potential pathogens, we focus only on the risk that farming and processing animals has on zoonotic disease emergence.

## 2. Methods

### 2.1. Approaching intensive chicken farming from an LCA perspective

We assessed peer-reviewed literature and industry documents detailing typical practices in intensive chicken meat production, with a key focus on identifying at which stages zoonotic pathogen spillover risks are present, and how the risks may vary across these stages. The primary source we used for information on intensive chicken production is the United States Department of Agriculture (USDA) “Poultry Industry Manual Foreign Animal Disease Preparedness and Response Plan”, which provides a detailed description of the stages involved in industrial chicken meat production (USDA, 2013). A graphical representation of this system described by (USDA, 2013) is shown as part of Figure 1. The main stages of intensive chicken production are primary breeding, parent breeding, hatching, grow-out, and processing. In the primary breeding stage, elite pedigree chickens are used to produce the parent breeding flock of chickens. The parent breeding stage is characterized by large houses of adult chickens (male and female), where mating and fertilized egg laying occur. Eggs laid at the parent breeding stage are transported to a hatchery, where they are incubated for roughly 20 days prior to egg hatching. Day-old chicks are then transported to grow-out houses, where they are fed for roughly 7 to 9 weeks. Finally, the 7–9-week-old chickens are transported to processing facilities where they are killed, de-feathered, and processed for sale. When their productive capacity decreases (usually after roughly 45 weeks of fertilized egg production), parent breeder chickens are also sent to the processing facility for slaughter and meat processing. Inputs to the system include feed, water, medicine/vaccines, litter, transportation, energy, and cleaning agents. Outputs of the system include meat and other items consumed by humans or other animals, as well as wastes like used litter, unhatched/broken eggs, dead or culled chickens during the production stages, feathers, blood, and wastewater. Used litter can be recycled (to some extent) in additional cycles of the grow-out phase of chickens (USDA, 2013), used as supplement for cattle feed (Charnas et al., 2025), or applied as agricultural soil amendment/fertilizer (Agga et al., 2025; USDA, 2013). Disposal of chickens that die or are culled during the production process, as well as feathers remaining after slaughter, can be disposed of in various ways, such as landfilling, composting, or incineration (Tesfaye et al., 2017; USDA, 2013). Furthermore, new methods for valorization of sidestreams from chicken production are being investigated, such as the extraction of proteins from feathers for use in materials (Soon et al., 2023).

**Figure 1.**
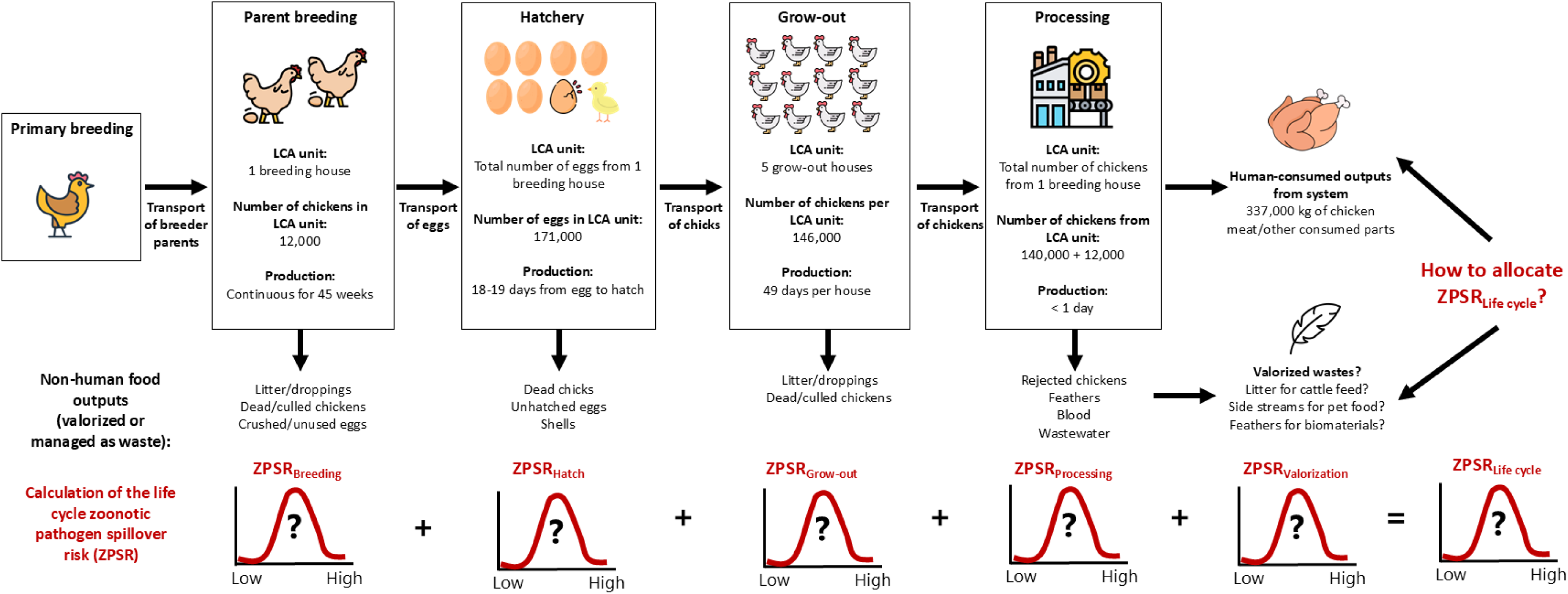
Conceptual overview of an intensive chicken production scheme (based on information in USDA 2013) and how the life cycle zoonotic pathogen spillover risk (ZPSR) can be assessed across this production chain. In this study, we expand on a semi-quantitative way of estimating the ZPSR at the grow-out stage (i.e., the *ZPSR_Grow-out_*, see Section 2.2). Icons created by gravisio, dDara, Ina Mella, Freepik, IconBaandar, Eucalyp, Luvdat, and Irfansusanto20 via Flaticon.com

There are various routes for the introduction of viruses into chicken farms, including potentially infected wild animals (e.g., birds or rodents) entering the chicken house, or workers carrying pathogen-containing material on clothes/shoes into chicken houses (Bartlett et al., 2022; USDA, 2013; Vora et al., 2023). A key factor in zoonotic disease risk within chicken production is the implementation of “biosecurity” practices that are put in place to limit or prevent the introduction of these pathogens into the farmed chicken population (see Section 2.2.2 for more details).

We propose a novel metric for assessing the risk of zoonotic pathogen spillovers from animal agriculture production: the life cycle Zoonotic Pathogen Spillover Risk (ZPSR). In its simplest form, the life cycle ZPSR is calculated as the sum of the pathogen spillover risks from each stage of chicken production, normalized by the maximum possible spillover risk from this sum, as shown in Equation 1:

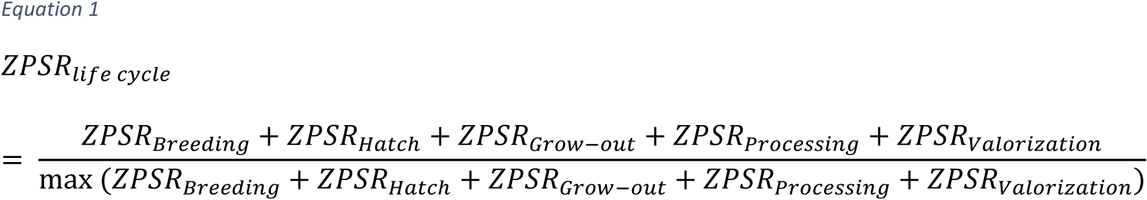

While the risks from each stage of chicken production are represented here as individual values, a more robust representation of these risks would be probability distributions of the zoonotic pathogen spillover occurring at each stage, as shown in Figure 1. We propose that the life cycle ZPSR be calculated based on the number of chickens/the amount of chicken meat and all processing sidestreams generated from one production cycle of one housing unit of a parent breeding facility (see Figure 1). We propose this unit of analysis starting from the parent breeding stage because the primary breeding stage has very high levels of biosecurity (USDA, 2013), thus decreasing the risk of pathogens entering the facility and spilling over into humans. Furthermore, for the parent breeding stage (and the grow-out stage), we focus on the zoonotic spillover risk at the level of the chicken house, as this is the unit at which key spillover processes operate (i.e., entry of a virus from wild animals into the chicken house, spread through the population in the chicken house, and transfer to workers in the chicken house). In developing estimates for the number of farmed animals at each stage of chicken production under this proposed life cycle approach, we primarily utilize methods and values described in agricultural guidance documents promulgated by the USDA (USDA, 2013) and university agricultural extension services (Brothers, 2022; Kester, 2025).

The number of fertilized eggs produced by the parent breeders over the course of their roughly 45 weeks of productivity (*n_eggs_*), as well as the number of chickens available for the grow-out stage (*n_grow-out_), can be calculated using Equation 2 and Equation 3:*

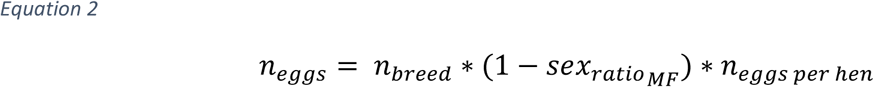

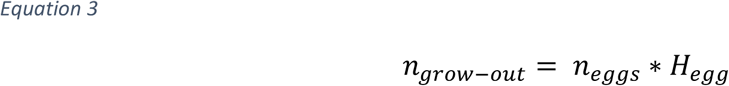

Where *n_breed_* is the number of parent breeding chickens in one typical breeding house (roughly 12,000; (French, 2017)), *sex_ratioMF_* is the typical ratio of male to female chickens (roughly 1:10), *n_eggs per hen_* is the average number of viable eggs laid per female in the duration of the breeding stage (157), and *H_egg_* is the fraction of eggs that successfully hatch (0.85). From this, the estimated number of eggs sent to the hatchery stage from one production cycle of a parent breeder house with 12,000 chickens is roughly 171,000 eggs, and the number of chickens produced for the grow-out stage from these eggs is 146,000. Given a grow-out stage housing size of 3,300 square meters and a housing density of 9 chickens/m^2^ (Brothers, 2022; USDA, 2013), this number of chickens would populate approximately 5 grow-out houses. With a loss rate of 3.6% of chickens during the grow-out stage (USDA 2013), the number of chickens sent to the processing stage would then be roughly 140,000, plus the 12,000 parent breeders when their productivity decreases. From this 152,000 chickens, assuming a weight at slaughtering of 2.95 kg and a dressing percentage of 75% (Kester, 2025), this would produce approximately 337,000 kg of chicken carcass weight for sale. While this mass of chicken would serve as the effective functional unit for calculating the life cycle ZPSR for chicken meat consumed by humans, for wastes/sidestreams from the chicken production process that are valorized via alternative uses (e.g., feathers), a different functional unit would be needed when assessing the ZPSR. Furthermore, details regarding how to allocate the life cycle ZPSR to different utilized outputs of the chicken production system (those consumed by humans or not) could take several approaches, whether it be on e.g., on a mass or economic value basis. This decision would likely be driven by various aspects of the specific products under consideration (e.g., the use of mass as the attribution metric if utilizing feathers from chicken production would likely not be an appropriate metric). We leave detailed discussion of allocation of the life cycle ZPSR for future work.

In the following section, we focus on developing a semi-quantification of the probability distribution for the zoonotic pathogen spillover risk for the grow-out stage of chicken production (i.e., the ZPSR_Grow-out_). We hypothesize that the ZPSR for this stage is among the highest of the different stages of chicken production. According to (USDA, 2013), the grow-out stage has the weakest biosecurity practices out of the other stages of chicken production, and also has a larger number of chickens and thus chicken houses (i.e., the unit at which spillover risk is considered in the grow-out stage) than the parent breeding stage. Furthermore, the hatching stage allows for less than a day of activity between hatched chicks before transfer to the grow-out houses, the processing stage provides little to no time for contamination with a wild-borne pathogen and spread through the flock, and the zoonotic risk from handling of wastes or sidestreams involves only contamination of material, and not introduction and spread of a virus among living birds. However, the stage-specific ZPSR probability distributions for these other stages should be prioritized for semi-quantification in future studies to enable a full assessment of the life cycle ZPSR (see Section 3.2 for more details).

### 2.2. Estimating zoonotic pathogen spillover risk at the grow-out stage

To semi-quantify the ZPSR_Grow-out_, we treat a spillover event as an “accident” and apply a modified version of existing methods around assessing the risk of an accident occurring across the life cycle of a product (i.e., (Khakzad et al., 2017), who develop an LCA of accident risk within oil and gas production). This involves developing an “event tree” of a series of events that must occur for a zoonotic pathogen spillover to happen (Figure 2). For a zoonotic pathogen spillover to occur in the grow-out phase of intensive chicken production, the hypothesized series of events required to occur are Event 1.) A pathogen must be present in wild animals in the area of the chicken farm, Event 2.) The pathogen must enter the chicken farm, Event 3.) The pathogen must infect and spread through the farmed chickens, Event 4.) The genome of the pathogen may have to mutate or reassort such that humans can be infected by it or be more easily infected by it, and Event 5.) The farm workers must become exposed to and infected with the pathogen.

**Figure 2.**
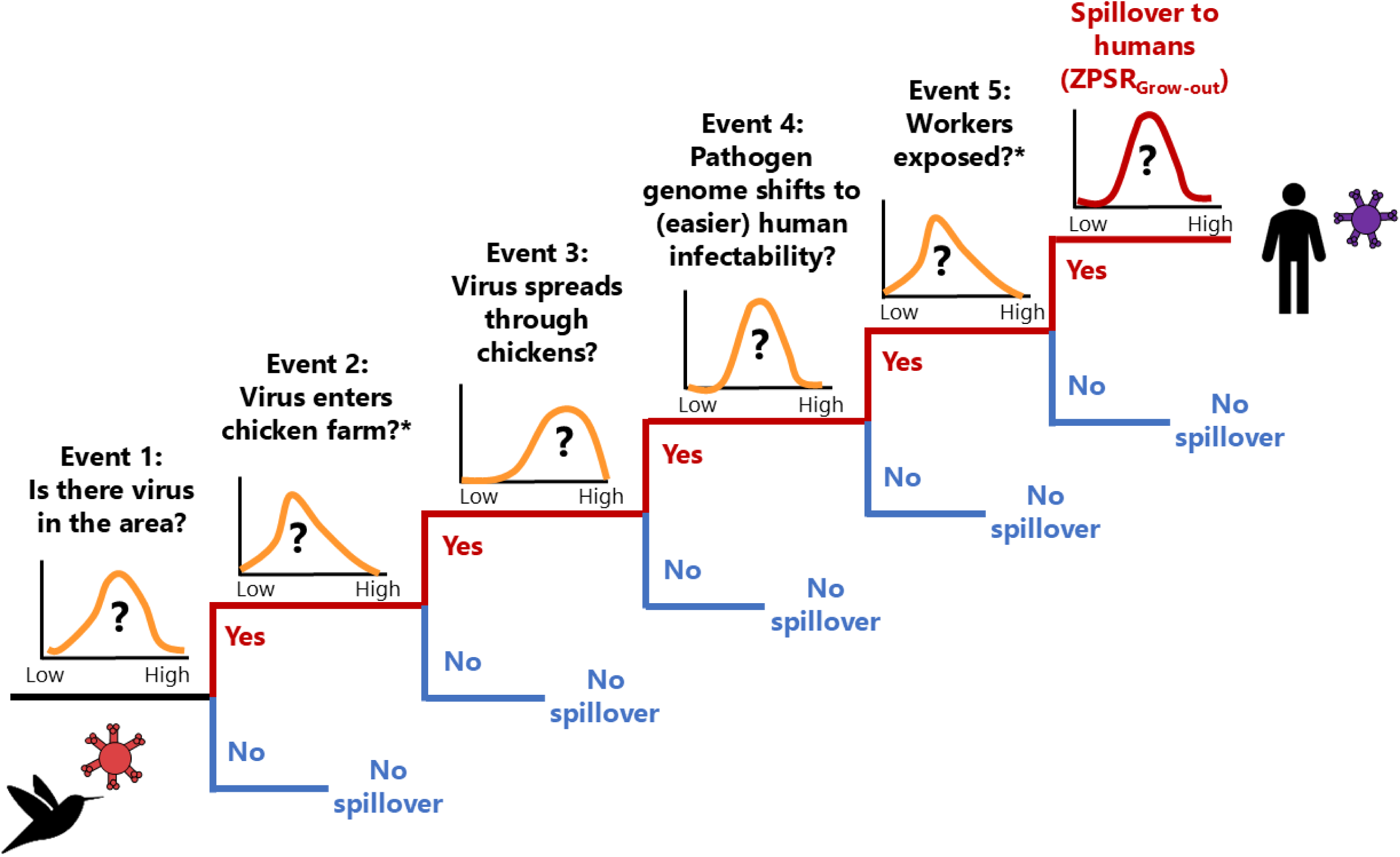
Event tree for a zoonotic pathogen spilling over from wildlife into humans during the “grow-out” stage of industrial chicken meat production, with illustrative probability distributions shown for each event. The final event’s probability distribution corresponds to the Zoonotic Pathogen Spillover Risk for the grow-out stage (ZPSR_Grow-out_). Note that Events 2 and 5 employ the same distribution (see Section 2.2.5).

The following sub-sections detail the approach taken to semi-quantify the likelihood of occurrence of each event in the event tree for a zoonotic pathogen spillover in the grow-out phase of intensive chicken production, focusing on the United States and Brazil. These are two of the largest chicken producing countries in the world (producing roughly 35% of chicken meat globally; (USDA, 2025)) and share many similarities in chicken production practices (Nääs et al., 2015; USDA, 2013). The goal with this quantification is to develop a distribution of risk values for each event, which would enable Monte Carlo analysis to ultimately arrive at a distribution for the likelihood of the final pathogen spillover event in the grow-out phase. As shown in Figure 2, the distribution of risk of each event would be scaled between being much-less likely (0 on the x-axis) to much more likely (100 on the x-axis) to occur. Table 1 provides an overview of how the risk distribution for each event in the event tree is semi-quantified, with further details provided in the sub-sections below. Unless otherwise noted, computational analysis was conducted using the R programming language 4.4.1 (R Core Team, 2024).

**Table 1.** Parameters used to semi-quantify the risk distributions at each event in the event tree for the grow-out stage of chicken production. HPAI = highly pathogenic avian influenza. SD = standard deviation.

| Event number | Event description | Data used to inform event risk distribution | Country-specific data used?<sup>1</sup> | Risk distribution defined by: | Data source(s) |
| --- | --- | --- | --- | --- | --- |
| Event 1 | Is there virus in the area of the chicken farm? | Number of chickens raised under intensive production, and number of wild bird species with migratory patterns overlapping chicken production regions.<br>Geospatial analysis performed on a 1° latitude/longitude grid. | Yes | Number of migratory bird species per grid cell, with sampling weighted by the magnitude of intensively raised chickens per grid cell | Number of intensively farmed chickens: (BirdLife International and Handbook of the Birds of the World, 2023)<br>Wild bird species ranges: (Gilbert et al., 2015)<br>Geospatial analysis: See Section 2.2.1 |
| Event 2 | Does the virus enter the chicken farm? | Data from surveys on the adherence to biosecurity practices at industrial chicken farms. | Yes | Details provided in Table 2 |  |
| Event 3 | Does the virus infect and spread through the chicken population? | Data from retrospective modelling on the "introduction time" of HPAI into a farmed chicken population, providing a distribution for the number of days between virus introduction and "detection time". | No | Risk distribution defined by number of days in the grow-out stage (49) and the risk of spread based on day of virus introduction:<br>Days 1-29: 100%<br>Day 35: 50%<br>Day 37: 5%<br>Days 37 - 49: 0% | Values extracted from Figure 1 of (Hobbelen et al., 2020) |
| Event 4 | Does the virus mutate to be able to (more easily) infect humans? | Literature values for the mutation rate of influenza A viruses (substitutions per nucleotide per cell infection) | No | Mutation rates of 7.1E-6, 3.1E-5, 3.9E-5, and 4.5E-5.<br>Mean = 3.1E-5, SD = 1.7E-5, cutoff values = 0 & (4.5E-5 + 7.1E-6) | From Table 1 of (Sanjuán and Domingo-Calap, 2016) |
| Event 5 | Are workers exposed to the virus? | Same risk distribution as Event 2 | Yes | Same risk distribution as Event 2 |  |

#### 2.2.1. Event 1: Is there virus in wild animals in the area of the chicken farm?

Wild birds, particularly migratory waterfowl, are important reservoirs for HPAI (Llanos-Soto et al., 2025; McDuie et al., 2024; Stiles et al., 2024). For this reason, we take the presence of migratory birds in a region where chicken farming occurs as a proxy for the possible presence of an HPAI virus in the area. We obtained data from Bird Life International on the geospatial ranges of >11,000 wild bird species globally, with identifiers for whether the birds are present in regions permanently, or on a migratory basis (see Figure 3, panels A & D). Additionally, we obtained data on the geospatial distribution of intensively farmed chickens globally from (Gilbert et al., 2015), who provide estimates for the number of intensively farmed chickens at 5 arc-minute grid spacing (Figure 3, panels B & E).

**Figure 3.**
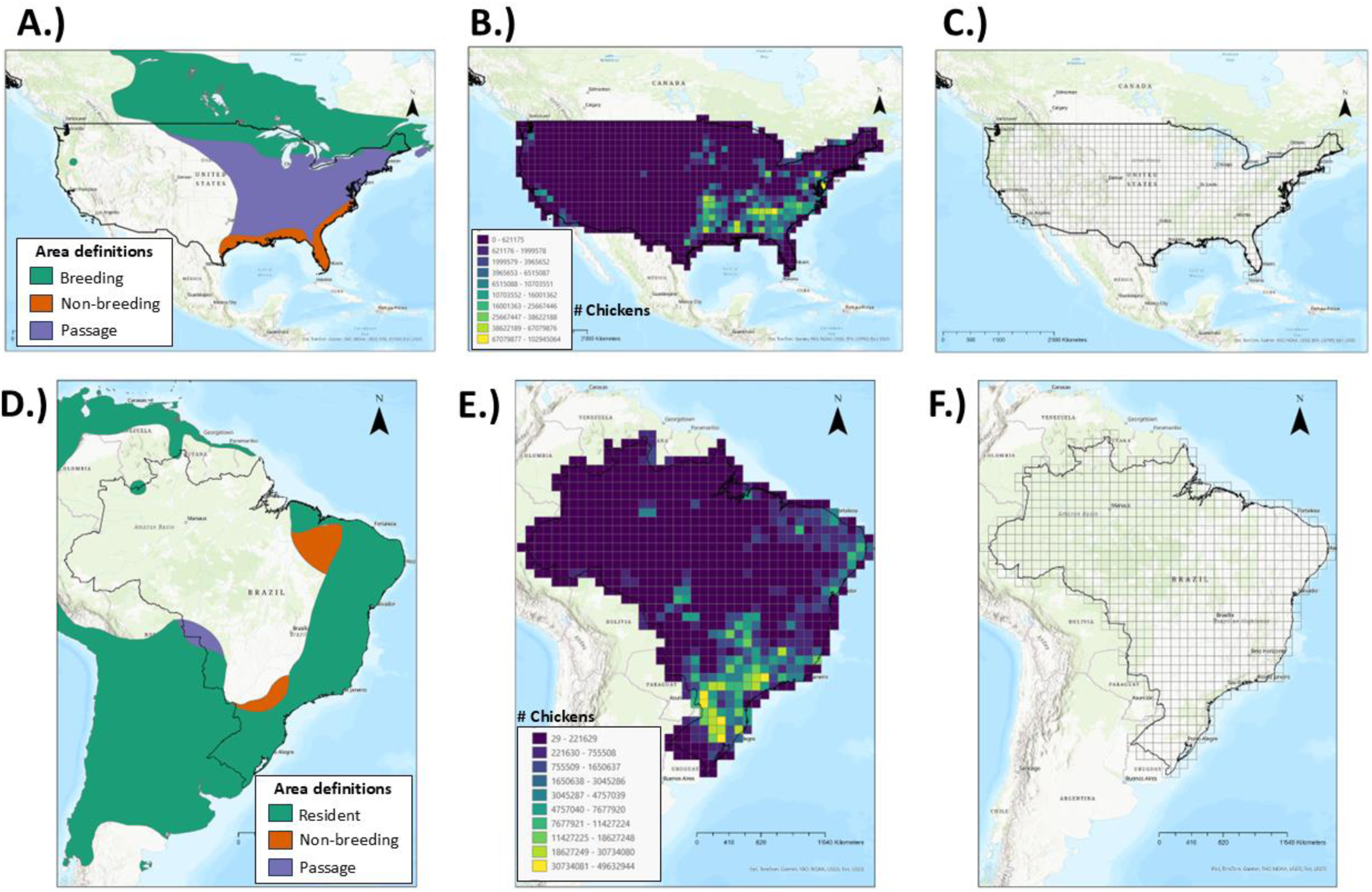
Panels A and D: Example geospatial migratory ranges for two bird species in the United States (US) and Brazil (data from BirdLife International and Handbook of the Birds of the World (2023)). Panels B and E: Estimated number of intensively farmed chickens on a 1° grid in the US and Brazil (data from Gilbert et al., 2015). Panels C and F: 1° latitude/longitude grids applied to the US and Brazil, on which the migratory bird and farmed chicken distributions are calculated.

Using ArcGIS Pro (Version 3.4: Esri), for both the contiguous United States and Brazil, we overlaid a grid with 1° latitude/longitude spacing (Figure 3, panels C & F). In each grid cell, we counted the number of migratory bird species whose migratory ranges (i.e., for breeding season, non-breeding seasons, or passage through a region) intersected with the analysis grid cell, and further normalized by the area (in km^2^) covered by the 1° grid cell to account for latitudinal differences in 1° grid cell areas. Projections used for calculating area were the USA Contiguous Albers Equal Area Conic projection for the contiguous US and the South America Albers Equal Area Conic for Brazil.

These numbers of migratory bird species per square kilometer in each 1° analysis grid cell were scaled to “relative risk” values between 0 and 100, normalizing between the US and Brazil using Equation 4:

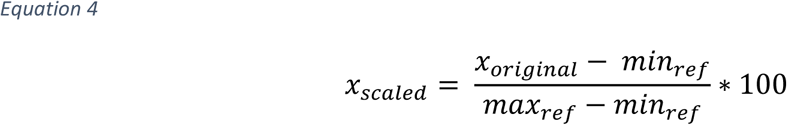

Where *x_original_* is the original number of migratory bird species in a grid cell, *min_ref_* is the reference minimum number of bird species possible (0 km^-2^), and *max_ref_* is the reference maximum number of bird species between either the US or Brazil (0.03 km^-2^). The resulting scaled and normalized values were used as the distribution of risk for the presence of virus in the area of chicken farming. Additionally, we summed the estimated number of farmed chickens in the 1° grid cells based on the data from (Gilbert et al., 2015), and normalized by grid cell area. We use this data on the number of farmed chickens per square kilometer in each 1° grid cell as a weighting for sampling the corresponding values of migratory bird species risk in the Monte Carlo modelling (see Section 2.2.6).

#### 2.2.2. Event 2: Does the virus enter the chicken farm?

Pathogens from wild animals can enter a chicken farming operation via several different routes. Wild birds can be attracted to the food and shelter available within a chicken farm, wherein they can spread pathogens to the farmed chickens. Rodents and insects can also be attracted to the conditions in chicken houses and be a vector for pathogen spread (USDA, 2013). Various human activities can also bring virus-containing material from outside of the farm to inside, given that HPAI can survive for up several days to several weeks depending on the environmental conditions (Kurmi et al., 2013). For example, workers on the chicken farm can inadvertently bring in pathogen-containing material from outside of the farm via e.g., their boots, tools, or wheels of their vehicles, providing a possible route of exposure for the farmed chickens (Dorea et al., 2010; Gkrinia et al., 2025; Islam et al., 2024).

The key determining factor in whether a pathogen from wild animals enters into a chicken farming operation is the level at which these different possible routes of pathogen introduction are mitigated, through so-called biosecurity practices. Biosecurity practices can differ between countries, provinces/states within a country, specific farming approaches (e.g., small-scale versus large-scale or enclosed versus free-range), and between contracting organizations that hire individual chicken farmers (De Oliveira Sidinei et al., 2021; Huang et al., 2017; Souillard et al., 2024). We identified two studies in the US and Brazil that conducted surveys of the actual on-farm implementation of biosecurity practices among chicken farmers in these countries, and used these to develop an initial estimation of the distribution of how strictly biosecurity practices are followed in the two countries of interest.

Briefly, we took statistics on the quantitative survey results in both countries and used these to develop distributions for how strictly the surveyed biosecurity practices were followed by the sampled population of chicken farmers. For Brazil, (De Oliveira Sidinei et al., 2021) surveyed 62 Brazilian chicken farmers in the state of Paraná, and assessed the adherence to 16 different biosecurity practices, including control of hand hygiene and footwear, a disinfection gate, and prevention of contact of the farmed chickens with wild animals (see Table 2). The farmers were ranked on their adherence to biosecurity practices on a scale from 0 (completely inadequate) to 10 (completely adequate). For each survey question, the mean and standard deviation of these survey results were reported across the surveyed population, with values broken down by farms with low biosecurity and those with high biosecurity (see Table 2). We took the weighted average of the mean values across the 16 biosecurity question results and across the two groupings of farms, as well as the weighted average of the standard deviation values, and used these to define an overall mean and standard deviation for biosecurity practice adherence for chicken farmers in Brazil. We generated a normal distribution of biosecurity practice adherence using this overall mean and standard deviation (using 1000 samples). Furthermore, the distributions of adherence to biosecurity practices were inverted to reflect that stronger adherence to biosecurity practices would result in lower risk of a virus from wild animals entering the chicken farm and removed values from the generated normal distribution that were below 0 and above 10.

**Table 2.** Details of risk distribution parameterization for the event of a pathogen entering the chicken farm (Event 2) and for workers being exposed to the pathogen (Event 5), as part of the zoonotic pathogen spillover risk event tree for the grow-out stage of chicken production (see also. **Figure 2). SD = standard deviation.**

| Country | Source of chicken farmer survey data | Geographic region & number of surveyed farmers | Details of survey data provided in sources | Data aggregation method for use of survey data in Event tree | Risk parameters |
| --- | --- | --- | --- | --- | --- |
| Brazil | Table 3 of (De Oliveira Sidinei et al., 2021) | State of state of Paraná<br>62 farmers | Ranked adherence to 16 biosecurity practices on a scale from 0 to 10.<br>Mean and SD of respondent rankings provided, aggregated to farms with low (19 farms) and high (43 farms) biosecurity. | Calculated weighted average of the mean and the standard deviation values across all biosecurity questions.<br>Weighting based on number of respondents in each group. | Risk distribution defined by:<br>Mean = 8.724, SD = 1.301<br>1000 Samples used to generate distribution in R (R Core Team, 2024). Cutoff values of 0 to 10.<br>Distribution inverted (to reflect worse biosecurity practices translating to higher risk) and re-scaled to 0 – 100. |
| United States | Table 3 of (Dorea et al., 2010) | State of Georgia<br>267 farmers | Frequency of adherence to 8 biosecurity practices were categorized as either "Never", "Rarely or Sometimes", "Usually or Always" followed.<br>Rankings provided for northern and southern regions of the state, and for visitors versus workers. | Calculated weighted average of "Never", "Rarely or Sometimes", and "Usually or always" adherence to each biosecurity practice.<br>Weighted based on number of respondents to each question.<br>Overall frequency of adoption: Never = 42.119, Rarely or Sometimes = 19.942, Usually or always = 37.939. | Frequency of adoption values used as weightings to generate risk distribution for viral entry, with:<br>"Usually or always" = 0% to 27% risk<br>"Rarely or sometimes" = 28% to 72% risk<br>"Never" = 73% to 100% risk |

For the United States, we utilized data from (Dorea et al., 2010), who provide results from biosecurity adherence surveys conducted on 267 chicken farmers in the US state of Georgia. Frequency of adherence to several biosecurity practices are reported as occurring “Never”, “Rarely or Sometimes”, or “Usually or Always”, with results separated for the two regions in the state surveyed (north versus south), and for visitors versus personnel at the farms (see Table 2). We used these data and the number of respondents to each question to calculate a weighted average across the survey questions for how frequently respondents adhered “Never”, “Rarely or sometimes”, or “Usually or Always” to the biosecurity practices. These weighted average values for biosecurity practice adherence were used to weight the generation of a distribution using 1000 samples, with the corresponding weighting value for the “Usually or Always” being used to fill viral entry risk values corresponding to 0 to 27%, the “Rarely or Sometimes” weighting being used to fill the risk values between 28 and 72%, and the “Never” category weighting being used to fill the top 27% of the risk distribution (see Table 2).

For both the US and Brazil, the risk distributions generated from the survey results were scaled to relative risk values using Equation 4, with *min_ref_* set to 0 and *max_ref_* set to 100. Overall, it should be noted that these biosecurity distributions should be taken as only preliminary examples for how information on biosecurity practices could be translated into distributions of risk that can be used in the zoonotic pathogen spillover event tree. We discuss the need for more comprehensive assessment of biosecurity risk distributions in our proposed LCA framework in Section 3.2.

#### 2.2.3. Event 3: Does the virus infect and spread through the chicken population?

To estimate the probability of a virus introduced into a chicken farm spreading through the population, we use estimates for the amount of time it takes between the introduction of an HPAI virus and when the infection in the population would be detected (detected being defined as when the observed mortality in the chicken population significantly exceeds that of the normal background mortality). To find this “introduction time”, (Hobbelen et al., 2020) took data from observed HPAI outbreaks in farmed bird populations, and used a “Susceptible, Exposed, Infected, Recovered” (SEIR) model paired with Monte Carlo simulations to probabilistically back-calculate values of the introduction time of the virus. We converted this probability distribution of their Monte Carlo simulations into a cumulative distribution function, with 5% of their simulations showing an introduction time of 12 days or less, 50% of simulations showing 14 days or less, and 95% of the simulations showing 18 days or less, with minimum and maximum values being roughly 12 and 20 days, respectively.

As shown in Figure 1, chickens are typically held in the grow-out stage for roughly 7 weeks (49 days) before transfer to the processing facility. Based on this and the results of Hobbelen et al., 2020, if an HPAI virus were introduced between day 1 and day 29 of the grow-out stage (i.e., with between 49 and 20 days left of the grow-out stage), there would be a 100% chance of the virus spreading through the chicken population enough to be detected. If the virus were introduced after day 29, the probability that the virus will spread through the population decreases to 50% by day 35 (i.e., 14 days left), to 5% by day 37 (i.e., 12 days left), and to 0% for days 37 to 49. These values were used to construct a distribution of risk values that the virus spreads through the chicken farm, assuming the likelihood of introduction into the chicken farm is equivalent at each day across the 49-day grow-out stage.

#### 2.2.4. Event 4: Does the virus mutate to be able to infect humans more easily?

For HPAI transmission to occur from birds to humans, close contact is generally required with a sick animal or contaminated material (ECDC, 2023; Miskiewicz et al., 2018). For currently circulating variants of HPAI, the viral genome is overall not well-adapted for infecting and transmitting between humans (Mehle et al., 2012; WHO, 2023). Thus, while evolution of the virus is not necessarily required for transmission to humans from farmed chickens now, additional evolution of the viral genome could make this more likely, thus increasing the risk of spillover events. Changes to the influenza viral genome occur via two distinct routes: genetic mutations and genetic reassortment (Mehle et al., 2012). Genetic mutations (i.e., errors) are introduced into copied RNA during viral particle replication. With genetic reassortment, two or more different variants of an influenza virus that are co-infecting the same cell of an infected organism exchange portions of genetic material. Changes in the virus’ capacity for infecting organisms (e.g., humans) via genetic mutations are relatively slower and smaller than they are through genetic reassortment. Furthermore, mutations that occur are random, and thus not deterministic towards more variants with more human infectibility. As a first-pass assessment towards semi-quantification for the risk of a variant that is more readily capable of infecting humans, we utilized information on the mutation rate of influenza viruses and discuss the limitations of this in Section 3.2.

We developed a proxy for a distribution of the risk of HPAI virus mutation resulting in a variant more capable of infecting humans. (Sanjuán and Domingo-Calap, 2016) provide values from 4 different studies on the mutation rate of influenza A viruses, specifically 7.1E-6, 3.1E-5, 3.9E-5, and 4.5E-5 substitutions per nucleotide per cell infection. We generated a normal distribution of virus mutation rates from the mean and standard deviation of these mutation rates (using 1000 samples), and removing sample values below 0, and above an artificial maximum of the highest plus the lowest reported mutation rates (i.e., 4.5E-5 + 7.1E-6). This distribution of mutation risk values was further scaled to relative risk values using Equation 4, with *min_ref_* set to 0 and *max_ref_* set to 100. This distribution was applied to both the US and Brazil.

#### 2.2.5. Event 5: Are the workers exposed to the virus?

Many of the biosecurity practices that would be followed on chicken farms, as described in (De Oliveira Sidinei et al., 2021; Dorea et al., 2010) and reflected in the Monte Carlo modelling of our study, would not only protect against the introduction of a virus into the chicken farm, but also against exposure of the workers to viruses. For this reason, we use the same risk distribution for the event of workers being exposed to the virus as we do for the likelihood of the event of a virus entering the chicken farm (see Section 2.2.2 for details).

#### 2.2.6. Monte Carlo modelling

Monte Carlo modelling was conducted using the distributions of relative risk values for Events 1 – 5 described in Sections 2.2.1 – 2.2.5. A series of 1E+3, 1E+4, 2.5E+4, 5E+4, and 1E+5 Monte Carlo simulations were conducted for both the US and Brazil risk distributions. Within each Monte Carlo simulation, one value of relative risk was randomly sampled from each event probability distribution, with the zoonotic pathogen spillover risk for that simulation being calculated using Equation 5:

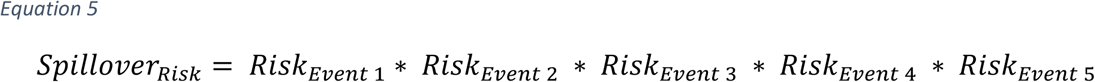

For the *Risk_event 1_* value, the sampling from the distribution of risk values (corresponding to the number of migratory bird species calculated per 1° grid cell) was weighted based on the corresponding number of intensively farmed chickens present in the 1° grid cells. This was done to ensure that areas where more intensive chicken farming was occurring would have the corresponding zoonotic risk from migratory bird species presence in that grid cell represented more in the overall spillover risk calculations.

## 3. Results and discussion

### 3.1. Zoonotic pathogen spillover risk in the grow-out phase

Figure 4 provides a scatterplot of the number of intensively farmed chickens per 1° analysis grid cell versus the number of migratory wild bird species, for the US and Brazil. There is large variability (roughly 6 orders of magnitude) in the number of farmed chickens/km^2^ in each grid cell in Brazil, for very little variability in the corresponding number of migratory wild bird species (between 0 and less than 0.01 species/km^2^). For the US, however, there is a much wider spread in the number of migratory bird species/km^2^ (between roughly 0.01 and 0.03) across a similar range of corresponding numbers of farmed chickens per grid cell as seen in Brazil.

**Figure 4.**
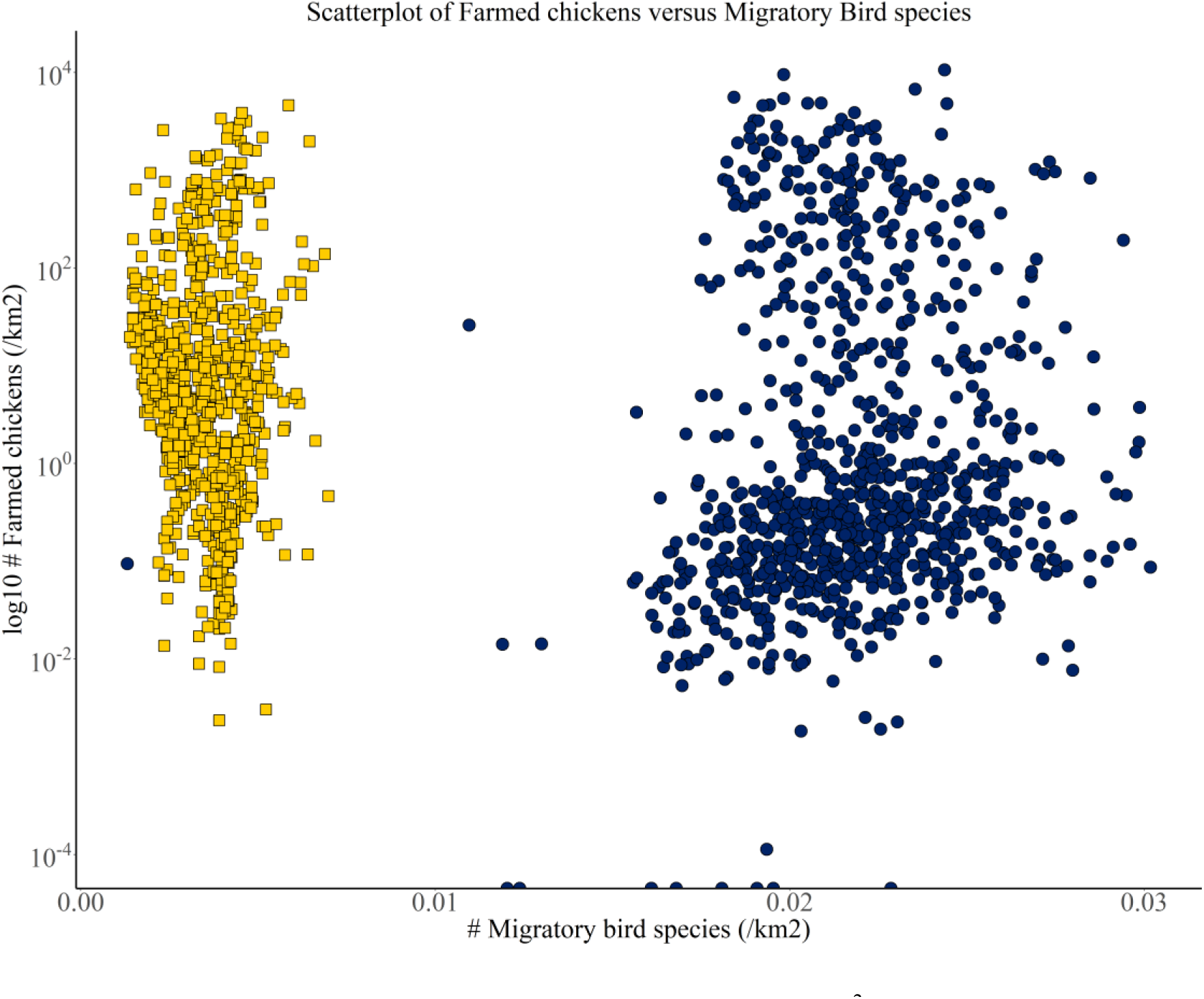
Scatterplot of the number of migratory bird species per km^2^ and the number of intensively farmed chickens per km^2^ for the 1° grid cells analysed over the US and Brazil. Values for Brazil are shown in yellow squares, while values for the US are shown in blue circles. The y-axis is log_10_ transformed.

Figure 5 shows the distributions of the relative risks of each event in the zoonotic pathogen spillover event tree. For the risk of virus being in the area of the chicken farm (driven by the presence of migratory wild bird species), both the US and Brazil show roughly normal distributions. However, in the US (as reflected in Figure 4), this distribution is much higher, centered over roughly 70 as opposed to roughly 12 in Brazil, indicating a much larger abundance of bird species with migratory patterns. Differences between wild bird migration patterns in North and South America are an area of active research in ornithology (Jahn et al., 2004; Meehan et al., 2022; Smith and et, 2022).

**Figure 5.**
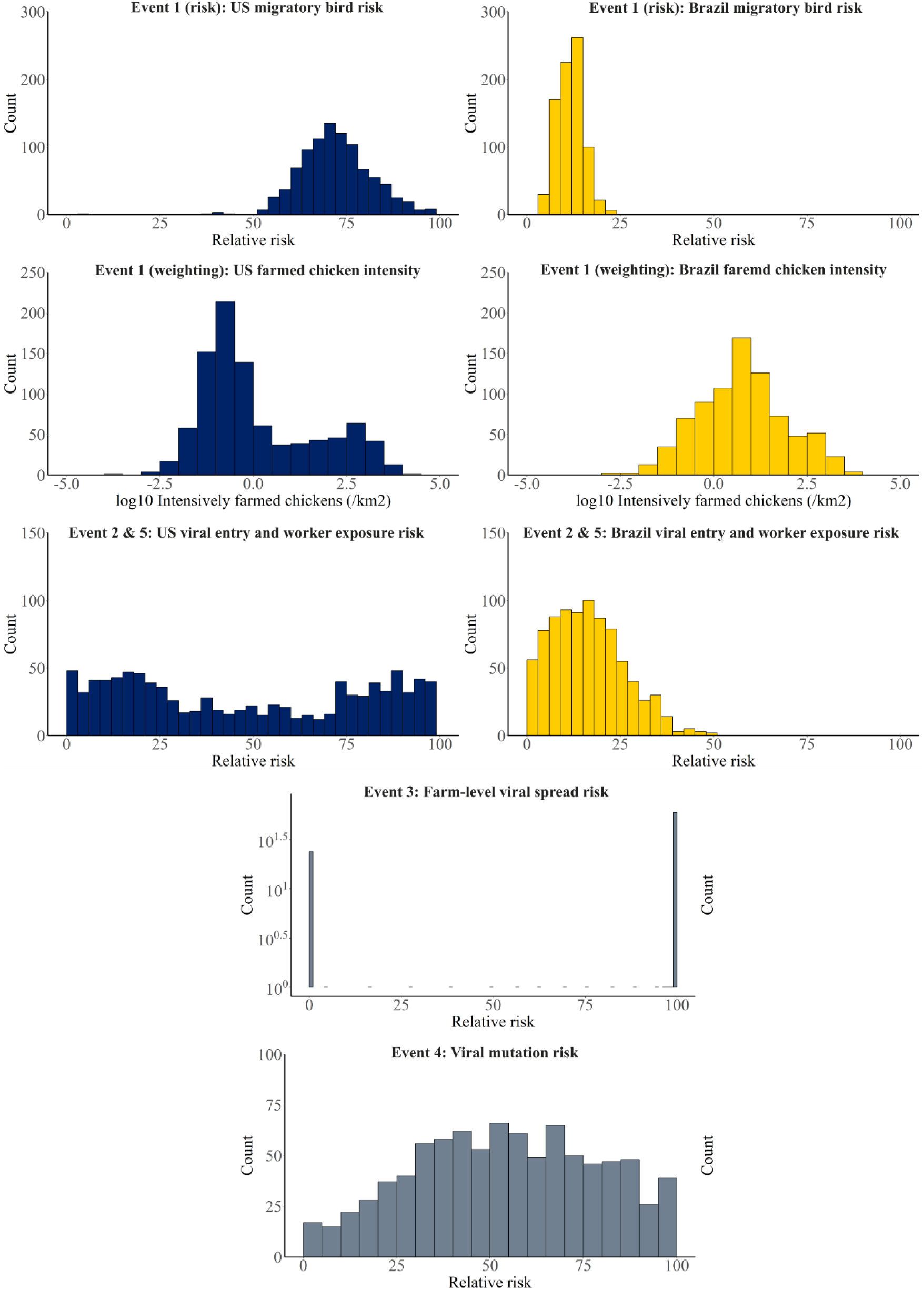
Distributions of the relative risks of events leading to a zoonotic pathogen spillover event during the grow-out stage of chicken production for chicken farms in the US and Brazil. Bar graphs shown in blue correspond to the US, while those in yellow correspond to Brazil. Bar graphs that are grey show identical distributions used for both the US and Brazil. Relative risk is defined as in Equation 4, and “Count” varies based on the total number of samples or observations used to develop the different risk distributions (see Section 2.2 for more details).

Regarding the risk of viral entry into the chicken farm, Figure 5 demonstrates that the stricter adherence to biosecurity practices reported by Brazilian chicken farmers results in an overall lower risk of viral entry compared to the US (recall that this same risk distribution is used for the worker exposure event, see Section 2.2.5). As mentioned in Section 2.2.2 though, we emphasize that these distributions should be taken only as preliminary, particularly since they are generated from just one study from each country on adherence to biosecurity practices. More robust and comprehensive analysis is needed to better understand large-scale differences between the biosecurity practices in the US and Brazil (see Section 3.2).

For the risk of viral spread occurring on the farm after the introduction of a virus, risk values are weighted heavily at either values of 100 or 0, reflecting introduction of the virus either well ahead of or well after when it is likely to have time to spread through the population (median “introduction time” of 14 days; see Section 2.2.3). Low frequencies of risk values between 0 and 100 correspond to the period of the grow-out phase (between days 29 and 37) during which the variability in “introduction time” assessed by (Hobbelen et al., 2020) would allow for potentially enough time for viral spread to occur.

Values for the relative risk for viral mutations (represented in our study as normally distributed) should also be interpreted with caution and effectively as a “placeholder” for more robust analysis around the risks of viral mutation and reassortment which could increase the risk of workers contracting the virus (see Section 3.2 for more details).

Figure 6 displays results for the estimated zoonotic pathogen spillover risk for the grow-out phase of intensive chicken farming, based on the Monte Carlo analysis. Regarding the number of simulations assessed, while the distribution in spillover risks demonstrates notably more smoothness going from 1E+3 to 1E+4 simulations (for the US results), little difference is seen between 1E+4 and 1E+5 simulations, suggesting 1E+5 simulations was an adequate number of simulations to capture the variability in all input parameters to the Monte Carlo analysis. Little to no variability is seen in the results for Brazil between 1E+3 and 1E+5 simulations.

**Figure 6.**
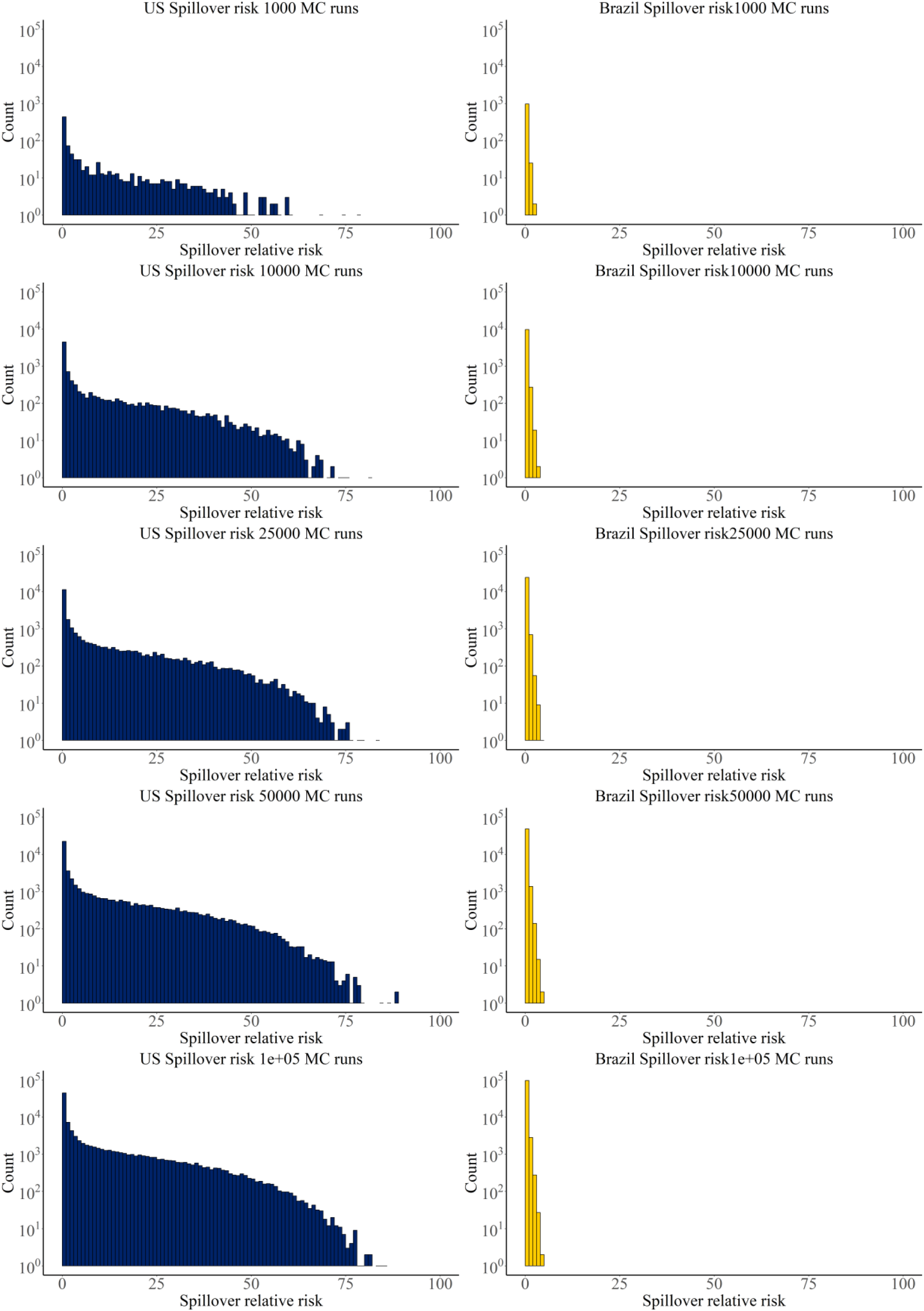
Overall zoonotic pathogen spillover risks for the grow-out phase of chicken production (ZPSR_Grow-out_) calculated from Monte Carlo simulations for the United States (left) and Brazil (right).

Overall, the US shows more frequent occurrences of higher risk values for a zoonotic pathogen spillover event than in Brazil. For example, in the analysis with 2.5E+4 simulations, all simulations in Brazil are below a 10% risk of resulting in a zoonotic pathogen spillover. In the US, while most simulations do have low risk, substantial risk is still present in a notable number of simulations (e.g., hundreds of simulations have spillover risk values around 50%).

These differential zoonotic pathogen spillover risk results between the US and Brazil should be interpreted with caution. The work we have done to semi-quantify the ZPSR_Grow-out_ probability distributions is a preliminary data gathering exercise, to understand what data is relevant and available for populating the event tree shown in Figure 2, and to enable a starting assessment of the ZPSR_Grow-out_ for two major chicken producing countries. The distributions of the relative risks of each event furthermore have large uncertainties and limitations (see Section 3.2). Despite these limitations, the overall result that the US has a larger background risk of HPAI introduction as a result of more migratory bird species being present aligns with climatological expectations. These findings also align with overall observations of the frequency of HPAI between the two countries, with the US having a substantial number of reported outbreaks of the ongoing HPAI H5N1 virus in commercial poultry farms, while Brazil has had few outbreaks reported (Cardenas et al., 2025; Martins-Filho and Quintans-Júnior, 2025; Mostafa et al., 2025).

Ultimately, these ZPSR_Grow-out_ probability distributions need to be built into the framework of the life cycle ZPSR. Since our event tree analysis for the grow-out stage utilized event probability distributions representative of the whole countries under analysis, we can map these results onto the framework discussed in Figure 1 by saying that the ZPSR_Grow-out_ probability distributions shown in Figure 6 correspond to the risk of a zoonotic pathogen spillover event occurring at one chicken house on a representative intensive chicken production farm in the US or Brazil. These risks should be scaled though to the number of grow-out houses in the LCA unit under analysis (i.e., 5 grow-out houses; see Figure 1). Additional research is necessary to expand the ZPSR probability distributions for the other stages of chicken production, as well as to develop appropriate scaling factors between these stage-specific ZPSR probability distributions to account for e.g., duration of the stage, the quantity of animals involved at each stage, and appropriate methods for allocation of life cycle ZPSR to different products (see Section 3.2).

### 3.2. Challenges and future research

We developed a framework for how to assess the life cycle risk of zoonotic pathogen spillover events in industrial chicken production, to be later fully embedded in operational LCA of animal agricultural production. We also developed a preliminary probabilistic semi-quantification of spillover risk for what is hypothesized to be the highest-risk stage of chicken production (the grow-out stage) for two of the world’s largest chicken producing countries, based around the “Event tree” model for accident risk (Khakzad et al., 2017). While this is a first step in expanding to larger-scale assessment of the life cycle risk of zoonotic pathogen spillovers from animal agriculture, there are several limitations with our study, and important areas of future work needed to embed this assessment framework into operational LCA practices.

A key limitation of our study is that the distributions of risk for virus entry into the farm and worker exposure were based on one study in each country of how strictly chicken farmers adhere to various biosecurity practices. For the US biosecurity survey information (Dorea et al., 2010), data are only from one state (Georgia), and the area of the state with the largest number of surveys collected from (the northern region, with 201 surveys compared to 66 from the southern region) had an active outbreak of infectious laryngotracheitis affecting chicken farms during the survey period, which could have biased the results of the survey (i.e., towards more strict biosecurity adherence). Similarly, for the survey data from Brazil (De Oliveira Sidinei et al., 2021), results were only from one state, and all respondents worked for the same contractor. Future studies could constrain these uncertainties by conducting a comprehensive assessment of the literature around biosecurity practice adherence across different countries. Additional data for on-farm implementation of biosecurity procedures at each life stage (e.g., vaccination, sanitation/protective equipment practices of human interactions, conditions during transport) should also be collected and made transparent to better understand how these factors vary spatially. Biosecurity assessment frameworks like the Biocheck.UGent tool (Gelaude et al., 2014) would be critical in standardizing this biosecurity information and making such information readily available to LCA modelers. Importantly though, while the Biocheck.Ugent website does provide country-aggregated data on surveys completed (Biocheck.UGent, 2025), no survey information is provided for the three highest broiler chicken producing countries in the world (the US, Brazil, and China; (USDA, 2025)). Furthermore, our approach to modelling the dynamics of pathogen spillovers from wild animal reservoirs to farmed chickens to humans takes several simplifying approaches regarding virus dynamics and biological interactions with hosts. This includes ignoring the impact of viral shedding rates from infected animals, environmental survival time of the virus, and dose/response relationships to exposed organisms (Plowright et al., 2017), which could be incorporated into future iterations of the LCA approach.

The key future research need in the context of advancing the LCA framework for zoonotic pathogen spillover risk is to semi-quantify the risks associated with the remaining steps in the chicken production value chain (i.e., parent breeding, hatching, processing, and valorization of wastes/sidestreams). For the parent breeding stage, the event tree would likely be structured in a similar manner, with the primary difference being that the overall biosecurity practices employed are higher than at the grow-out phase (USDA, 2013), and thus the risks of viral entry and worker exposure would likely be lower. For the hatching stage, live chicks spend only roughly one day post-hatching in these facilities (USDA, 2013), with therefore limited time for introduction of a virus into the farmed chicken population. However, the potential for pathogen entry and contamination into hatching facilities during the incubation period (roughly 18 days), and its potential dynamics with the chick population, should be explored further for assessment of this risk. Transmission of wild virus to the farmed chickens during transport between the hatching and grow-out stage and the grow-out stage to processing facility could also be a key contributor to zoonotic risk (Linder et al., 2023), with the dynamics of exposure likely to be very different than that during the breeding or grow-out stages. Contamination of trucks used between different farmed chicken flocks could be a unique route for pathogen introduction that is absent from the core parts of the production stages (Linder et al., 2023). Regarding the processing stage, close human contact with the slaughtered chickens and their derived parts are risk factors for zoonotic spillover (MacMahon et al., 2008), which would present substantially different risk factors than during the other stages. A better understanding of the risk distributions at these other stages, and the overall life cycle ZPSR, would also be beneficial for developing standardized methods for allocating the ZPSR to the different utilized outputs of intensive chicken production, whether it be human-consumed food or sidestreams. Furthermore, on a larger scale, the impact of mass culling operations of farmed chickens or other animal populations in response to infection with a zoonotic pathogen would also contribute to the overall environmental impacts of chicken production, due to e.g., decreased productivity (Capper, 2023; Gickel et al., 2025). The impacts of these disease events on farmed animal product life cycle environmental impacts should be fully incorporated into LCA approaches, in tandem with the risk of zoonotic pathogen spillovers that we propose here.

The risks of zoonotic pathogen spillover from the handling and use of sidestreams from the chicken production process (e.g., used litter, feathers, blood) are potential routes for pathogen transmission to humans (Yamamoto et al., 2010). While used chicken litter (containing the bedding litter, feathers, feces, etc.) can be used as agricultural fertilizer or feed addition for cattle (Agga et al., 2025; Charnas et al., 2025; USDA, 2013), and chicken feathers removed at the slaughtering and processing stage are often sent for incineration or landfilling (Tesfaye et al., 2017), the increasing interest in alternative uses for agricultural sidestreams in the “bioeconomy” could result in blood, feathers, and other animal processing sidestreams being diverted for use in, for example, extraction of keratin from feathers for protein-based advanced materials or the use of proteins extracted from animal blood in bioplastics (Ding et al., 2025; Soon et al., 2023). With these changing dynamics for the use and exposure to potentially pathogen-containing sidestream materials in alternative valorization routes, the zoonotic pathogen spillover risk dynamics of this stage of the chicken production value chain are crucial to better understand and incorporate into the overall LCA framework in the near future.

While this study focused on the life cycle zoonotic pathogen spillover risks associated with the on-farm risks during the chicken production life cycle, spillover risks would also exist from the growing of feed crops for farmed animals. In particular, regions that are recently deforested, and at agriculture/pristine wilderness buffer zones, may be at particularly high risk for zoonotic spillover events from wildlife into humans (Carlson et al., 2025). Such spillover risks would also need to be attributed to crops grown for human consumption on recently deforested land (Morand and Lajaunie, 2021). Close collaboration with researchers incorporating the impacts of agriculture/human activities on biodiversity into life cycle assessments (Cabernard et al., 2024) will be important for incorporating this land use aspect of zoonotic pathogen risk into LCA. However, the dynamics associated with land-use change and zoonotic spillover risk can be quite complex (Carlson et al., 2025) and likely do not lend themselves to the event-tree approach taken with on-farm spillover risks. Possible relationships between this zoonotic pathogen spillover risk and the increasing interest in incorporating animal welfare into LCA (Scherer et al., 2018; Turner et al., 2023) should also be explored within the LCA community.

Expanding this framework to assess the zoonotic spillover risks across the life cycle of farming other animals that are key potential “bridge species” for zoonotic spillovers from wildlife into humans (e.g., pigs) should also be prioritized by the LCA community. Some work has already been done on spatially-analyzing risk factors for spillover events of e.g., Nipah virus emergence in Thailand (Thanapongtharm et al., 2019) and influenza from backyard pig farming in Mexico (Mateus-Anzola et al., 2019). Quantitative approaches that exist for simulating virus spread considering animal population size (Etbaigha et al., 2018), transport of animals (Wongnak et al., 2020), and jumping of viruses to humans (Royce and Fu, 2020) could also be incorporated into the quantitative approaches for further developing the framework introduced here.

Furthermore, large environmental impacts can occur from viruses present in wild animals entering domesticated animal populations, amplifying in pathogenicity, and spilling back into the wild animal populations (Dhingra et al., 2018; Xie et al., 2023). This is another key environmental impact – in particular, to biodiversity – that animal farming practices can cause, which are currently unaccounted for in LCA frameworks of animal agricultural practices.

Additionally, a range of methods exist for translating various agricultural emissions or activities into human health or environmental effect metrics, such as translating life cycle CO_2_ emissions to the midpoint level of global warming potential, which is further translated to the endpoint level of impact on human or ecosystem health (Huijbregts et al., 2016). Translating the life cycle zoonotic pathogen spillover risk developed in this study into human health or ecosystem effect endpoints (such as Disability Adjusted Life Years; DALYs) would be a long-term goal of this methodology. However, key differences between the dynamics of zoonotic pathogens that spill over into humans and other impacts assessed from agriculture would complicate these human or environmental effects from being incorporated. While zoonotic disease spillover events can cause harm to the exposed individual (as has been seen e.g., in dozens of cases of HPAI virus spilling over from cattle or other farmed animals to farm workers in the US in 2024 – 2025; (CDC, 2025; Garg et al., 2024)), much greater impacts would be caused if the spillover event leads to further human-to-human transmission, an epidemic, or a pandemic. Such large-scale human health impacts from a zoonotic pathogen (possibly) having spilled over from agricultural animals into humans has occurred multiple times in recent decades (e.g., Swine flu and Middle East Respiratory Syndrome; (Rabaan et al., 2021; Shapshak et al., 2011)) but these epidemic/pandemic dynamics depend on many societal factors such as access to health care, policies, and societal infrastructure for detecting and containing emerging pathogens (UNEP, 2016). Future work in developing characterization factors towards this end could build on previous work that assesses geographic variability in zoonotic disease response readiness (e.g., (Sun et al., 2024)).

## 4. Conclusions

Increased awareness of the various facets of the environmental impacts of animal agriculture have impacted governmental policies and consumers’ choices, resulting in increasing action towards lowering the environmental impact of animal agriculture. Many of these aspects are currently incorporated into LCA and other sustainability assessments of animal agriculture, providing policymakers and consumers with information on how they can lower their environmental footprint through changes in diet. More awareness, transparency, and quantification of the impact of animal agriculture methods on the risk of zoonotic pathogen spillovers would provide stakeholders insight into a key area of the environmental impact of animal agriculture that is currently largely not covered by impact ranking systems. The novel LCA assessment framework we have developed in this study for zoonotic pathogen spillover risk in chicken production, and the semi-quantitative assessment of a key stage of production (the grow-out stage), are a first step towards a more complete and robust assessment of animal agriculture for comparison to other food sources. Furthermore, emerging approaches to the use of by-products from the agricultural system, such as chicken processing sidestreams for valorization into the bioeconomy, should be prioritized for quantification into this risk framework to ensure such environmentally conscious approaches are conducted in a truly sustainable manner.

To shift towards sustainable agricultural practices, our understanding of the impact of animal agriculture on zoonotic disease emergence should be incorporated in decision making, just as greenhouse gas emissions and land use impacts are. This approach would also support the ambitions of (Hobeika et al., 2023) who outline the potential need to develop an Intergovernmental Panel for One Health for pandemic prevention, preparedness, and response, as well as the World Health Organization’s current efforts around an international instrument on this topic (WHO, 2025).

## Funding and acknowledgements

This project has received funding from the ETH Board of Switzerland in the framework of the Joint Initiative: Proteins for a Sustainable Future. The authors thank Roland Hischier from Empa for his helpful discussions associated with this study

## CRediT authorship statement

**John D. Hader:** Conceptualization, methodology, investigation, formal analysis, software, visualization, writing – original draft. **Nadia Malinverno:** Conceptualization, writing – review and editing. **Claudia Som:** Conceptualization, funding acquisition, supervision, writing – review and editing.

## Data statement

All sources for data used in this study are provided in Tables 1 and 2. Data for intensively farmed chickens, biosecurity practice adherence in Brazil and the United States, viral spread risks in a farmed chicken population, and viral mutation rates are publicly available. The geospatial data for wild bird distributions are not publicly available, but can be requested from BirdLife International via the following weblink: http://datazone.birdlife.org/species/requestdis

